# Native, Spatiotemporal Profiling of the Global Human Regulome

**DOI:** 10.1101/2025.06.14.659727

**Authors:** Lindsay K. Pino, Daniele Canzani, Andrea Gutierrez, Julia Robbins, Brian McEllin, Evan Hubbard, Erin Broderick, Anastasiya Prymolenna, Lillian Tatka, J. Sebastian Paez, Gaelle Mercenne, Kyle Siebenthall, William E Fondrie, Alexander J. Federation

## Abstract

The regulome, comprising transcription factors, cofactors, chromatin remodelers, and other regulatory proteins, forms the core machinery by which cells interpret signals and execute gene expression programs. Despite its central role in development, disease, and drug response, the regulome remains largely uncharted at scale due to its dynamic, low-abundance, and chromatin-associated nature. Here, we present a method for scalable, regulome profiling for global, compartment-resolved quantification of native regulome proteins. By enriching DNA- and chromatin-associated proteins and profiling them using high-throughput, label-free DIA mass spectrometry, regulome profiling captures chromatin-associated proteins across 36 human cell lines and thousands of perturbations. The resulting Regulome Atlas recovers nearly 60% of known human transcription factors, reveals lineage-specific TF localization, and distinguishes active nuclear engagement from latent, unbound states. We demonstrate that regulome profiles resolve acute immune pathway activation prior to transcriptional changes, identify previously unrecognized drug-induced regulome responses, and enable proteome-scale readouts of compound target engagement and complex remodeling. This work establishes a foundational resource for decoding the regulatory proteome and provides a blueprint for integrating regulome data into next-generation models of cellular behavior.

## INTRODUCTION

New measurement technologies have consistently expanded our scope of biological understanding over the past 25 years. Each generation of assays from DNA sequencing to high-throughput proteomics and single-cell approaches has enabled access to previously unmeasurable types of biomolecules and biological information, revealing new layers of regulatory control and reshaping our understanding of cellular function. Increasingly, these datasets serve not only as reference maps, but as essential training material for artificial intelligence models that aim to reconstruct and predict cellular behavior ^1–3^. Building a functional virtual cell begins with the ability to comprehensively measure its molecular components ^4,5^.

Projects such as the Human Genome Project, ENCODE, and GTEx have characterized gene structure, regulatory elements, splicing, and interindividual variation in expression across hundreds of primary human tissues and cell types using many technologies ^6–10^. More recent efforts have extended these maps into single-cell and spatial dimensions ^11,12^, while others have developed high-resolution methods to pinpoint nucleosome positioning and factor occupancy at regulatory elements in situ ^13–15^. These epigenomic atlases have been instrumental in identifying enhancer landscapes, understanding lineage specification, and interpreting noncoding disease variants.

For the proteome, tissue- and context-specific atlases have mapped protein abundance, turnover, and interaction networks across developmental stages and organ systems^16–18^. Spatially resolved proteomic efforts have further refined our view of organelle organization and protein localization^19^. These resources have proven foundational for defining cell types, prioritizing disease biomarkers, and nominating drug targets. Despite these advances, a critical component of the cellular regulatory machinery remains poorly characterized, the ***regulome***.

The regulome is defined as the complex ensemble of proteins that integrate cellular signaling pathways and then bind DNA and chromatin to dynamically orchestrate transcriptional programs^20–22^. The regulome encompasses the complete ensemble of proteins that integrate cellular signaling pathways and orchestrate transcriptional programs through interaction with DNA and chromatin. This includes not only sequence-specific transcription factors, but also cofactors, chromatin remodelers, histone-modifying enzymes, RNA-binding regulators, and architectural proteins that collectively determine gene expression outcomes. Critically, regulome activity depends not just on protein abundance, but on nuclear localization, post-translational modifications, and dynamic assembly into functional complexes at chromatin^23,24^. These proteins play central roles in development, homeostasis, and disease, yet remain difficult to measure at scale due to their low abundance, transient binding behavior, and extensive post-translational regulation^22,25,26^.

While genomics-based assays such as ATAC-seq, DNase-seq, and motif analysis provide insights into protein localization on the genome ^27,28^, and ChIP-seq has enabled targeted profiling of individual factors, a comprehensive, and scalable atlas that quantifies the proteins that make up the regulome has remained out of reach. Even the ENCODE project, among the most expansive efforts in functional genomics, has directly profiled less than 3% of known chromatin-associated regulatory proteins by ChIP-seq at great effort and cost ^9^.

Here, we present a method to address this gap with scalable, regulome-wide protein profiling. By coupling efficient chromatin fractionation with high-throughput, label-free DIA mass spectrometry, this approach enables global, compartment-resolved quantification of native DNA- and chromatin-associated proteins, capturing key components of the regulome without genetic tagging or crosslinking^29,30^. We use this platform to construct the Regulome Atlas, a first-of-its-kind resource detailing the abundance and nuclear localization of transcription factors, cofactors, and other regulatory proteins across 36 human cell lines spanning major tissue types and cancer lineages. Beyond static maps, we apply the method to track regulome dynamics in response to cellular stimuli and small molecule perturbations. This work establishes a scalable framework for native, unbiased measurement of chromatin-bound proteins and provides a foundational dataset for studying the dynamics of the regulome.

## RESULTS

### A Scalable Regulome Workflow That Recapitulates Canonical Chromatin Biology

Building on the principles of chromatin enriching salt separation coupled to DIA (CHESS-DIA) ^31^, we have optimized extraction conditions for high-throughput enrichment of chromatin- and DNA-associated proteins for compatible with end-to-end processing in 96-well plates and magnetic bead handling robotic systems (**Figure 1A**). Specifically, we developed a lectin-conjugated magnetic bead that binds specifically to glycans enriched on the nuclear membrane. This lectin is more specific for the nucleus than Concanavilin A, which also binds strongly to cell membrane glycans (**Figure S1A**).

**FIGURE 1.**
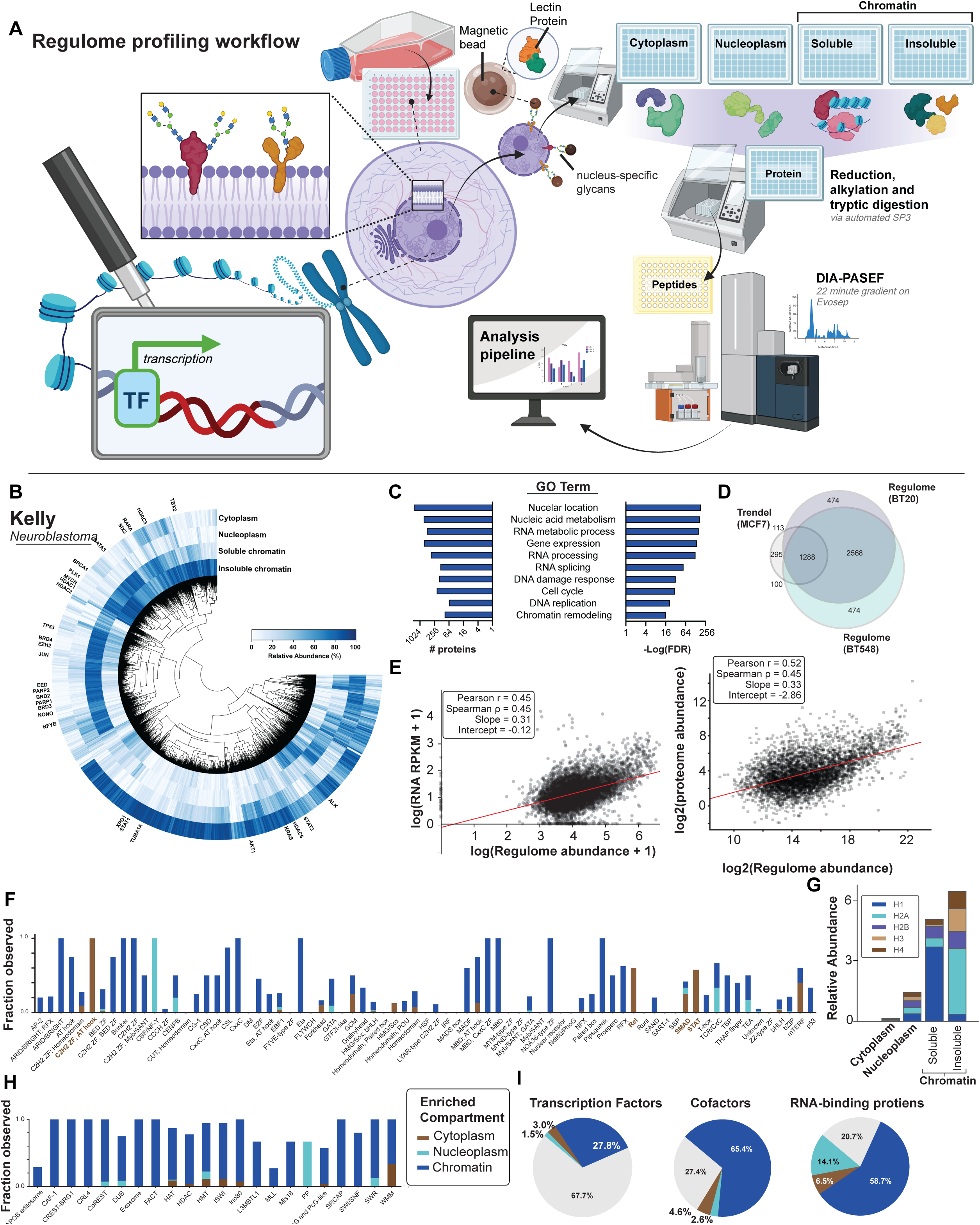
An Overview of the Regulome Profiling Platform and the Chromatin-Associated Proteome in Kelly Neuroblastoma Cells **(A)** Schematic of the regulome profiling platform workflow. Cells are fractionated into cytoplasmic, nucleoplasmic, and chromatin compartments through sequential enrichment steps. Proteins are prepared via automated SP3 processing and peptides are analyzed using mass spectrometry. **(B)** Circular histogram of the median-normalized protein abundances in each subcellular fraction of Kelly neuroblastoma cells. **(C)** Gene Ontology (GO) term enrichment for proteins identified in the chromatin fraction, derived from a representative cell line (neuroblastoma cell line Kelly). Enriched categories include chromatin-associated and RNA processing functions. **(D)** Comparison of shared and unique protein detections between a photo-crosslinking method^42^ for regulome enrichment in the breast cancer cell line MCF7 versus this proposed method in BT20 and BT548. **(E)** Correlation of regulome measurements using this proposed method protein abundance versus RNA expression (left, Pearson r =0.45) and whole-cell proteome abundance (right, Pearson r = 0.52). **(F)** Detection frequency of transcription factor DNA-binding domain families across the dataset. Proteins of each DBD family were assigned to the dominant fraction in which they were observed. **(G)** Relative abundance of histone proteins, grouped by histone type, across fractions. **(H)** Enrichment of protein complexes. Frequency of chromatin-related complex components (e.g., SWI/SNF, HDAC, MLL, CoREST) across compartments highlights the platform’s capacity to recover functional transcriptional machinery. **(I)** Classification of detected regulome proteins. Pie charts indicate the distribution of TFs, cofactors, and RNA-binding proteins, with 67.7% localized to chromatin.

Cells are dissociated, washed, followed by isolation of cytoplasm leaving intact nuclei. The nuclei are then manipulated using magnetic lectin-cinjugated beads and exposed to a minimal set of extraction buffers to separate proteins in the nucleoplasm from the chromatin-bound proteins. Subcellular protein fractions are digested into peptides with SP3 method^32^. We then apply label-free quantitative data independent acquisition mass spectrometry (LFQ-DIA-MS) methods to detect and quantify the resulting peptides^33,34^. The resulting data, for a given cell line sample, consists of the unbiased proteome quantification in each of the four resulting subcellular proteome fractions (cytoplasm, nucleoplasm, soluble chromatin and insoluble chromatin) (**Figure 1B**).

Focusing on the chromatin fraction, these proteins are enriched for Gene Ontology (GO) terms associated with markers of expected DNA-binding functions like nuclear localization, gene expression, and DNA damage response; additionally, these proteins are associated with RNA-focused functions such as RNA metabolic process, RNA processing, and splicing (**Figure 1C**). Notably, the abundance of chromatin-bound regulome proteins correlates, weakly with corresponding RNA transcript abundances (r = 0.45) and whole-proteome abundances (r = 0.52. **Figure 1E**) ^35^, a similar level of correlation that is typically observed between RNA and global protein levels. The shared variance (R² = 0.20-0.27) demonstrates that approximately 75-80% of the biological variation captured by regulome profiling represents unique regulatory protein states not measured by standard RNA-seq or whole-cell proteomics approaches (**Figure S2A**). Shannon entropy analysis of this dataset revealed high information content (2.58 ± 1.48 bits per protein) with clear biological discrimination the cell lines measured, reflecting the dynamic range and condition-specific regulation accessible through chromatin enrichment. These correlations are consistent with previous studies showing that protein localization and activity are poorly predicted by transcript or total protein abundance alone ^36–38^. The uncorrelated variance captured by regulome measurements likely represents biologically meaningful information about nuclear import, post-translational modifications, protein complex formation, and competitive DNA binding that determines functional TF activity.^39–41^.

This accessible and scalable approach to enriching for regulome proteins associated is in concordance with other regulome measurement approaches, with high overlap between this biochemical fractionation method and a recently described photo-crosslinking method (**Figure 1D**)^42^, despite the different breast cancer cell lines profiled in the two studies. Of all the DNA-interacting proteins detected by photo-crosslinking (1805 proteins in MCF7), we captured ∼70% (1118 proteins of the 1805 Trendel et al proteins detected in all four breast cancer cell types assessed with regulome profiling), with an additional 2500+500 unique proteins detected here. The high degree of overlap across different breast cancer cell lines validates the shared core regulome components that bind directly to DNA, while the non-overlapping proteins likely reflect both cell-type-specific regulatory programs and the complementary nature of these approaches. Photo-crosslinking captures proteins in direct DNA contact at the moment of UV exposure, while regulome profiling measures the broader population of chromatin-associated regulatory machinery, including cofactors and chromatin-modifying complexes that may not directly contact DNA but are functionally engaged at chromatin. The method presented here also correlates highly with other previously reported methods for chromatin purification^31^ (**Figure S1**).

We first examined the distribution of histones across fractions. Linker histone H1, which binds DNA more transiently and is released under milder extraction conditions, is enriched in the soluble pool. In contrast, core nucleosomal histones (H2A, H2B, H3, H4) remain predominantly in the insoluble pellet, likely due to their tight association with the DNA in the nucleosome, reflecting the biochemical accessibility of chromatin-associated proteins under our fractionation conditions (**Figure 1G**)^31^.

When considering DNA-binding TFs as annotated by DNA binding domain (DBD)^43^, we find the vast majority of TFs are found bound to chromatin, although some families such as the signaling responsive TF families like STAT, SMAD and REL (NF-kB) are predominantly identified outside the nucleus (**Figure 1F**). Finally, when considering protein members associated with epigenetic regulators which physically interact with chromatin rather than directly with DNA^44^, we again find that the majority of these complexes are measured in the chromatin fraction (**Figure 1H**), with an exception of a three-protein histone H2AX phosphatase complex (PP)^45^ that was detected primarily in the cytoplasm for this cell type and condition.

In data from Kelly neuroblastoma cells, we detect 32.3% of annotated TFs, 72.6% of cofactors, and 79.3% of RNA-binding proteins. Of the proteins detected, the majority are localized to the chromatin compartment (**Figure 1I**). Of note, the majority of chromatin associated proteins are not captured in these three categories, which account for 25% of the detected chromatin-associated proteins. In the remaining 75% of the chromatin proteome, we see expected categories including histones, DNA replication machinery, protein homeostasis proteins (ubiquitin and deubiquitin proteins), and protein translation. Interestingly, nearly 30% of the observed chromatin proteome have no annotated Gene Ontology molecular function classification (**Figure S1**). Together, these data highlight the diversity and complexity of the chromatin-associated proteome while pointing to a vast unexplored space within the regulome.

### A Regulome Atlas of chromatin-bound proteins across cell types

To observe how regulome composition varies across tissue types, we built the first atlas of chromatin-bound active human regulomes by profiling a panel of 36 human cell lines. This panel spans a broad range of adult and pediatric tumors including carcinomas, sarcomas, hematologic malignancies, and central nervous system cancers, with representation across both common and historically understudied indications (**Figure 2A**).

**Figure 2.** Landscapes of the Chromatin-Bound Regulome Across Cancer Cell Lines. **(A)** Overview of the 36 human cell lines, spanning diverse tissue origins including carcinomas, sarcomas, hematologic malignancies, and CNS tumors, with representation from both adult and pediatric cancers. **(B)** Cumulative detection of 897 unique transcription factors (TFs), representing ∼60% of the annotated human TF repertoire, across the full cell line panel. **(C)** Histogram showing the number of cell lines in which each TF was detected, revealing that nearly half of TFs were observed in ≤3 cell lines, while a smaller subset shows widespread chromatin occupancy. **(D)** Comparison of TF coverage and per-sample efficiency per cell line in detecting TFs between regulome profiling and large-scale proteomics datasets (CCLE, CPTAC, DrugMap, GTEx, Knol, PROCAN). **(E)** As in D, comparison of TF coverage and TF detection efficiency normalized to instrument time required to construct the dataset. **(F)** Heatmap showing relative abundance of detected RNA-binding proteins, chromatin regulators, and TFs across all 36 cell lines. Rows represent proteins; columns are samples. **(G)** UMAP projection of TF abundance across cell lines, with canonical master regulators colored by their lineage-specific identities (e.g. AR in prostate, GATA3 in breast, PHOX2B in neuroblastoma). **(H)** Dot plot of select regulome protein abundances on chromatin across selected cell lines, grouped by indication. Dot size corresponds to median-normalized abundance.

Despite the relatively small number of samples, we detected 897 unique annotated TFs, nearly 60% of the annotated human TF repertoire (**Figure 2B**). 47% of TFs were restricted to 3 or fewer cell lines, with 15% of TFs showing broad expression across all cells in the atlas (**Figure 2C**). Despite the more technically complex cellular fractionation methods, the extent of TF coverage rivals or exceeds that of large-scale proteomics efforts such as CCLE, CPTAC, DrugMAP, GTEx, TPCPA, and PROCAN^10,35,46–49^ (**Figure 2D**). Notably, we achieve this depth using a fraction of the instrument time and many fewer samples. The streamlined workflow captures significantly more TFs per hour and per sample than these broader atlases (**Figure 2E**).

The Regulome Atlas recovers many TFs known to act as master regulators of development and key drivers of lineage-specific cancers (**Figure 2F, G**). Androgen receptor (AR) is selectively engaged in prostate cancer lines, while FOXA1 is shared in both prostate and breast cancers; MYOD1 and PAX3-FOXO1 mark rhabdomyosarcoma (Gryder et al. 2019); PHOX2B and HAND2 collaborate in the core regulatory circuit of MYCN–amplified neuroblastoma, while GATA3 also plays a role in neuroblastoma along with specific subtypes of breast cancer ^50^(Durbin et al. 2022); and Brachyury (TBXT) is uniquely active in chordoma (Chen et al. 2020). These patterns hold across diverse contexts with HNF4A in hepatocyte lines, PAX8 in ovarian cancer, and OTX2 in medulloblastoma all consistently localized to chromatin. STAT3, typically cytoplasmic, shows strong and specific chromatin engagement in a subset of cell types associated with response to STAT3 inhibition, highlighting a potential regulatory shift not evident from total protein abundance alone^51,52^.

Integrating the Regulome Atlas with ATAC-seq data shows that TF abundance helps resolve common ambiguities in motif analysis. Motifs for TFs with high chromatin occupancy are more strongly enriched in accessible regions, suggesting that TF localization, not just motif presence, shapes chromatin accessibility. This becomes especially important when multiple TFs share similar motifs. For example, in prostate cancer, the FOXA motif is enriched, but regulome data confirms that FOXA1, not FOXA2, is the dominant chromatin-bound factor in PC3 cells (**Figure S3**). Similarly, while HOXA motifs are broadly enriched, only specific paralogs are active in a given context; HOXA2, for instance, has been linked to a regulome switch driving resistance to androgen deprivation therapy^53^.

Together, these data illustrate how regulome profiling not only recapitulates known transcriptional programs but also reveals unexpected nuclear activity, offering a dynamic view into the logic of lineage identity and disease state.

### Spatiotemporal regulome dynamics during immunological stimulation

Next, we sought to observe the acute regulome dynamics only visible through time-respolved spatial proteomics measurements. For this, we stimulated various immune pathways and observed regulome changes in a single cell type. We analyzed THP-1 monocytic leukemia cells which are commonly used as a surrogate for innate immune response in a monocyte context^54^. 14 conditions were assessed at two physiologically relevant doses, whose known effects span TLRs 1-9, NOD1, NOD2 and the STING pathway. THP-1 cells were treated for 2 hours to capture early, proximal changes in protein localization prior to major transcriptional or chromatin remodeling events, then processed for regulome profiling (**Figure 3A**).

**Figure 3.**
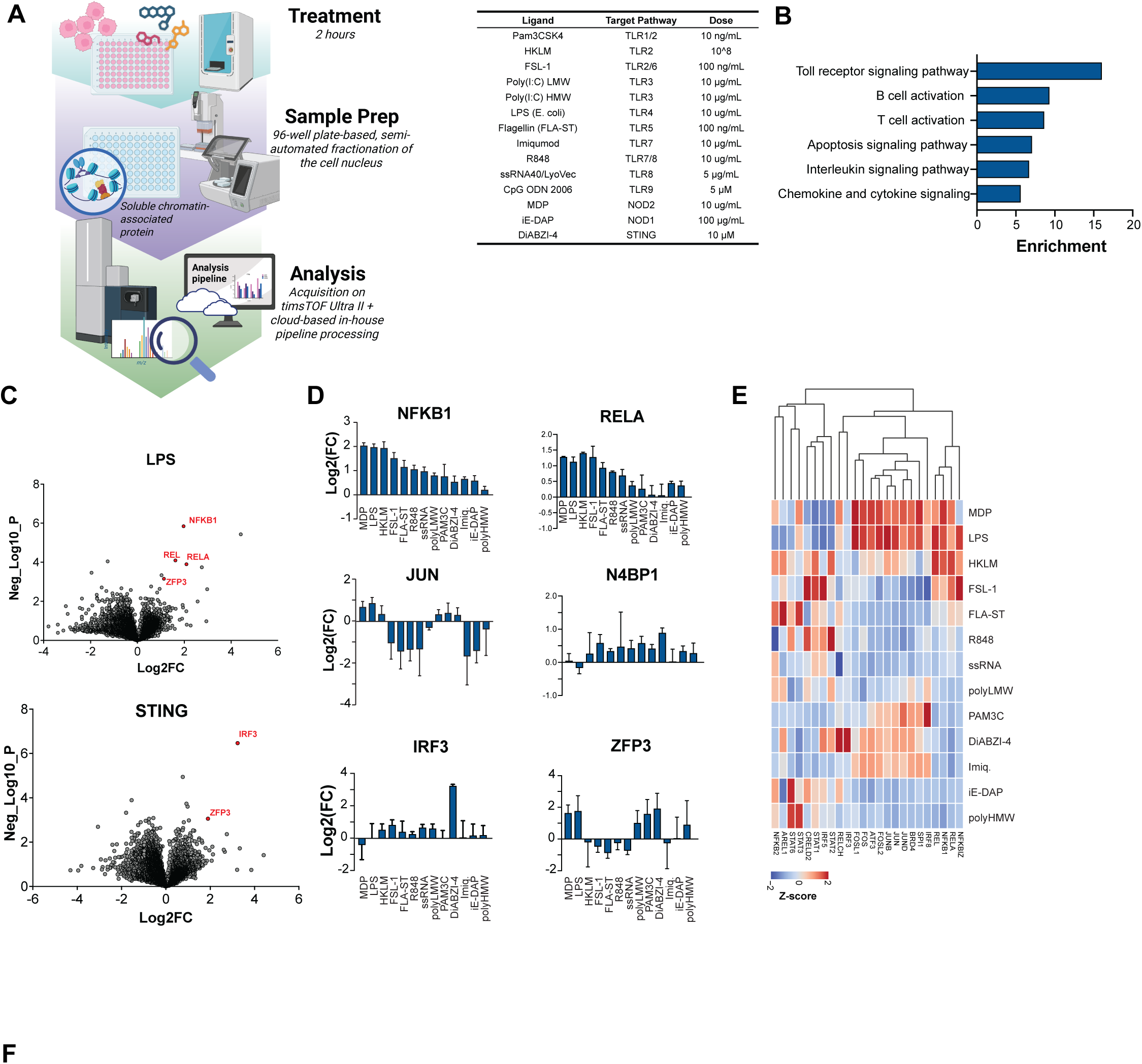
Chromatin-Associated Protein Profiling of Immune-Stimulated THP-1 Cells (A) Experimental workflow for immune stimulation assay. THP-1 monocytic leukemia cells were treated with 14 innate immune ligands (table inset) targeting TLR1–9, NOD1, NOD2, and STING signaling axes. Cells were harvested after 2 hours and processed in 96-well plate format for chromatin enrichment and DIA-MS analysis. (B) Gene Ontology enrichment analysis of proteins showing increased chromatin association post-stimulation. Enrichment scores are computed across all 14 conditions. (C) Volcano plots of differential chromatin-associated protein abundance following treatment with LPS (top) or STING ligand DiABZI-4 (bottom), where log2 fold change compared to DMSO is plotted against negative log10-transformed p-values for each protein. (D) Log2 fold change in chromatin occupancy for selected transcription factors, cofactors, and signaling proteins across all stimulation conditions. Each row represents a protein; columns correspond to specific ligands. (E) Z-score normalized abundance heatmap of selected proteins detected in the chromatin fraction across all immune stimuli. Data reflect relative signal intensity from DIA-MS.

Regulomes were analyzed to identify proteins significantly changing chromatin occupancy after stimulation. Proteins increasing abundance on chromatin were highly enriched for GO terms related to known pathways including TLR signaling, B and T cell signaling, interleukin signaling and chemokine signaling (**Figure 3B**)^55^. All conditions showed significant increase in chromatin occupancy of the NFKB1 TF, as expected (**Figure 3D**).

MDP-1, a dipeptide motif sensed by cytosolic NOD2, activates NF-κB via RIPK2 and TAK1 but lacks access to IRF or AP-1 axes and does not amplify through feedforward loops. As a result, MDP-1 induces a minimal chromatin response, with strong NFKB1 recruitment and only 18 additional chromatin-associated proteins, reflecting a highly specific transcriptional program for early antibacterial defense^56^. By contrast, FSL-1 (TLR2/6 ligand) and HKLM (heat-killed Listeria) engage multiple TLRs, triggering MyD88/TRIF pathways and activating NF-κB, MAPKs, and IRFs. These stimuli recruit hundreds of chromatin-associated proteins (301 for FSL-1, 263 for HKLM), consistent with a broad transcriptional response integrating multiple signaling arms (**Figure S4**)^57^.

Canonical immune activators LPS (TLR4) and STING ligand DiABZI-4 recapitulate known TF activation: NFKB1, RELA, and REL are enriched in the LPS condition; IRF3 dominates the STING response^58^. Key coactivators (e.g., p300, CREBBP, BRD4) and kinases (e.g., CHUK, IKBKE) show stimulus-specific chromatin recruitment patterns aligned with their canonical signaling roles (**Figures 3D, 3E, S4**).

Curiously, a poorly characterized zinc finger protein, ZFP3, showed consistent chromatin enrichment across multiple distinct stimuli, including LPS, STING, MDP-1, and PAM3C, not the broad effect seen with HKLM (**Figure 3D**). While its precise mechanism remains unknown, the consistent recruitment pattern, particularly alongside master regulators like NFKB1 and IRF3, positions ZFP3 as a high-priority candidate for functional characterization. This finding exemplifies how unbiased regulome profiling can identify previously overlooked regulatory factors that may play important roles in immune signaling.

### Remodeling of the chromatin-associated proteome after drug treatment

Given the varied regulome effects observed in response to immune stimuli, we moved to measure the acute effects of small molecule drugs on the regulome. To accomplish this, we profiled 299 select compounds that span the unique mechanisms of action seen in FDA-approved drugs (**Figure 4A, Supplementary Table 3**). Compounds were tested in triplicate, leading to 4.36 million observations of compound:protein effects across this experiment, yet required only 15 days of mass spectrometry acquisition time. Data quality was high, with median DMSO CV across the experiment of 14% and high correlation within and across plates (average r^2^=0.962 intraplate, **Figure S5**).

**Figure 4.**
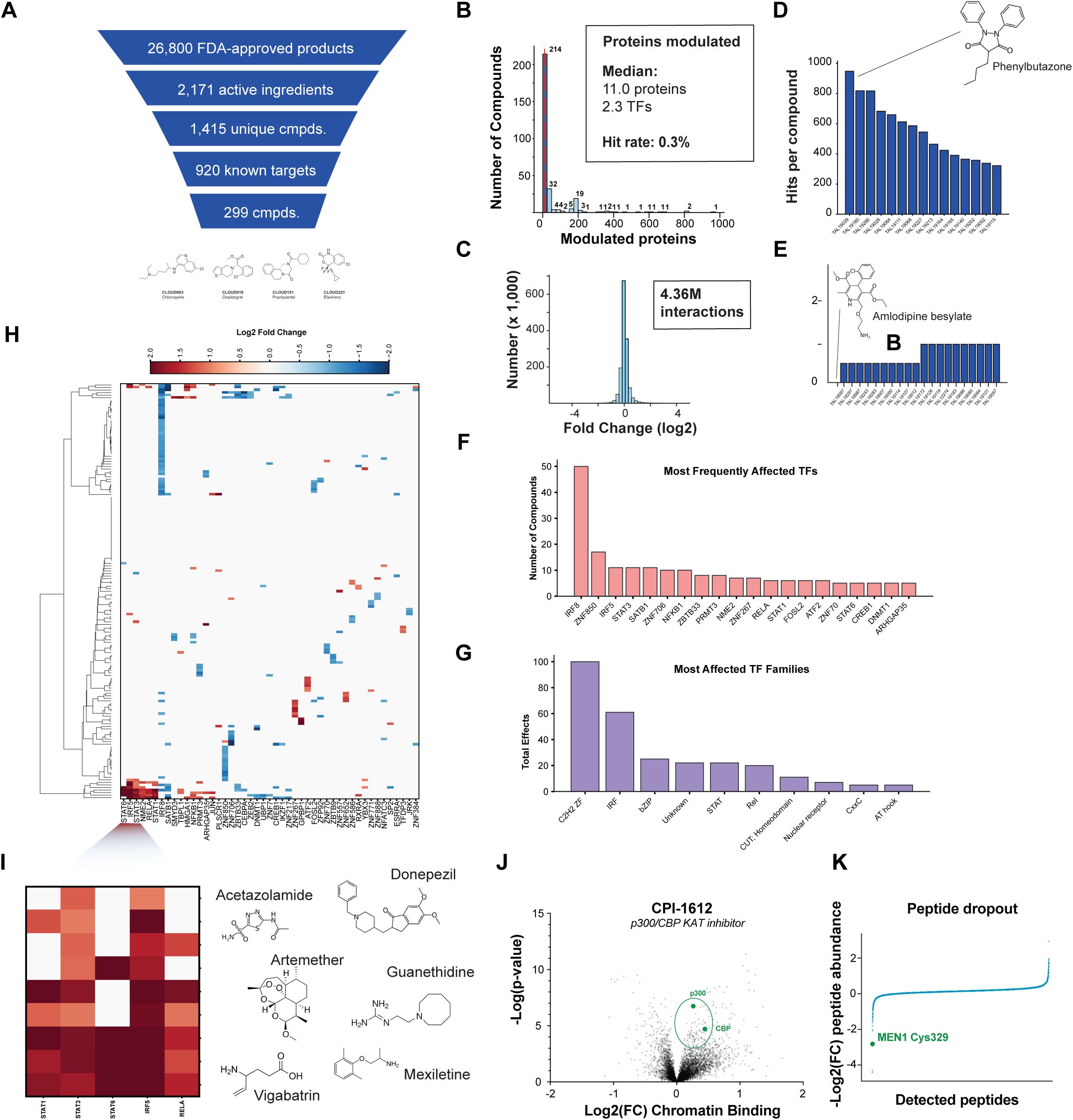
Regulome response to a diverse library of bioactive small molecules (A) Schematic overview of the curation pipeline used to assemble a set of 299 small molecules with diverse annotated mechanisms of action. (B) Histogram showing the number of significantly modulated proteins per compound (|log₂FC| > 1, BH-adjusted p < 0.05). (C) Distribution of all compound–protein interactions in the dataset, shown as log₂ fold changes. (D–E) Bar plots highlighting compounds with the most (D) and fewest (E) protein-level changes. (F) Transcription factors with the highest number of distinct compound-induced modulations. (G) TF families most frequently modulated, grouped by DNA-binding domain annotations from *Lambert et al*. (H) Heatmap of transcription factor responses across the compound library, restricted to TFs with the highest modulation rates. Non-significant effects are shown in grey. (I) Cluster of compounds that activate at least two of five immune-associated TFs: STAT1, STAT3, STAT6, IRF5, or NFKB1. (D) (J) Volcano plot showing protein-level changes following treatment with CPI-1612 (1 µM, 24 h), a p300/CBP inhibitor. (E) (K) Waterfall plot showing global protein abundance changes (median-normalized peptide intensity) after treatment with M-1121 (1 µM, 4 h).

Compound effects against regulome proteins were calculated, using a minimum threshold of 2-fold change and adjusted p-value < 0.05 across the dataset. With these parameters, we observed 8,646 instances of decreased regulome abundance and 8,552 instances of increased abundance, representing a rate of 0.3% for observed protein modulation. The median compound caused an effect on 11.0 proteins and 2.3 TFs (**Figure 4B, C**). Highly-active compounds caused changes in hundreds of proteins. For example the nonsteroidal anti-inflammatory drug (NSAID) Phenylbutazone resulted in 642 significant protein depletions and 176 protein increases, consistent with its high observed clinical toxicity and removal from the market (**Figure 4D**)^59^. Many compounds resulted in no or few effects, like the calcium channel blocker Amlodipine besylate showing zero significant changes to the regulome, for example (**Figure 4E**).

461 TFs showed modulation from at least one compound, with several TFs and TF families showing frequent modulation, especially those related to inflammatory pathways (**Figure 4F, G**). This is consistent with the important role that monocytes play an early responder in the innate immune response^60^. IRF8 was the most modulated TF, along with IRF5, STAT3 and NFKB1.

We identified a discrete cluster of compounds (acetazolamide, artemether, donepezil, guanethidine, vigabatrin, and mexiletine) that consistently induced immune-related TFs (**Figure 4H, I)**. Despite acting on unrelated primary targets, all six compounds converged on a shared regulome signature marked by strong induction of inflammatory TFs, including RELA (NF-κB p65), STAT3, STAT5, and IRF5. This suggests that diverse cellular perturbations can trigger common transcriptional stress responses, possibly reflecting a convergent inflammatory or innate immune activation state. While the precise upstream pathways vary, prior studies have linked many of these compounds to cellular stressors such as ROS generation, lysosomal disruption, or metabolic imbalance, each of which can activate NF-κB or JAK–STAT signaling cascades under certain contexts. However, further work is required to dissect the molecular drivers of this shared TF profile ^61–63^.

In addition to the functional effects on TFs observed across the compound library, we also were able to observe signs of compound:target engagement as well. For example, an enzymatic inhibitor of the CBP/p300 histone acetyltransferase domain led to increased abundance of both CBP and p300 protein levels on chromatin, possibly due to stabilization of a chromatin-bound protein conformation (**Figure 4J**). For covalent compounds, we can also observe specific loss of peptides containing cysteine sites where the compound is irreversibly bound, due to the resulting change in the peptide’s mass. For example, we see M-1121, a covalent MLL:Menin inhibitor leading to decreased abundance of the C329-containing peptide, while other peptides in the Menin protein show attenuated loss (**Figure 4K**).

### MLL complex dynamics

Observing the robust target engagement of M-1121, we next dissected how small molecules that disrupt the Menin:MLL protein-protein interaction impact the MLL1-containing COMPASS complex. We performed regulome-scale proteomic profiling in dose-response in THP-1 cells, a model of MLL-AF9–driven leukemia that retains key molecular features of oncogenic rewiring. In these cells, MLL fusion proteins sustain aberrant transcriptional programs, including activation of HOXA genes and MEIS1.

Treatment with the covalent inhibitor M-1121 led to broader and more potent remodeling of chromatin-associated complex composition than the non-covalent VTP50469, suggesting distinct biochemical consequences of Menin engagement (**Figure 5A**). Regulome quantification revealed chromatin displacement of Menin and its accumulation in the nucleoplasm—consistent with redistribution rather than degradation (cytoplasm not analyzed). Despite Menin displacement, MLL1 (KMT2A) remained chromatin-bound across all doses, indicating that disruption of the MLL:Menin interaction is not sufficient to fully disassemble the MLL1 complex (**Figure 5B,C**).

**Figure 5.**
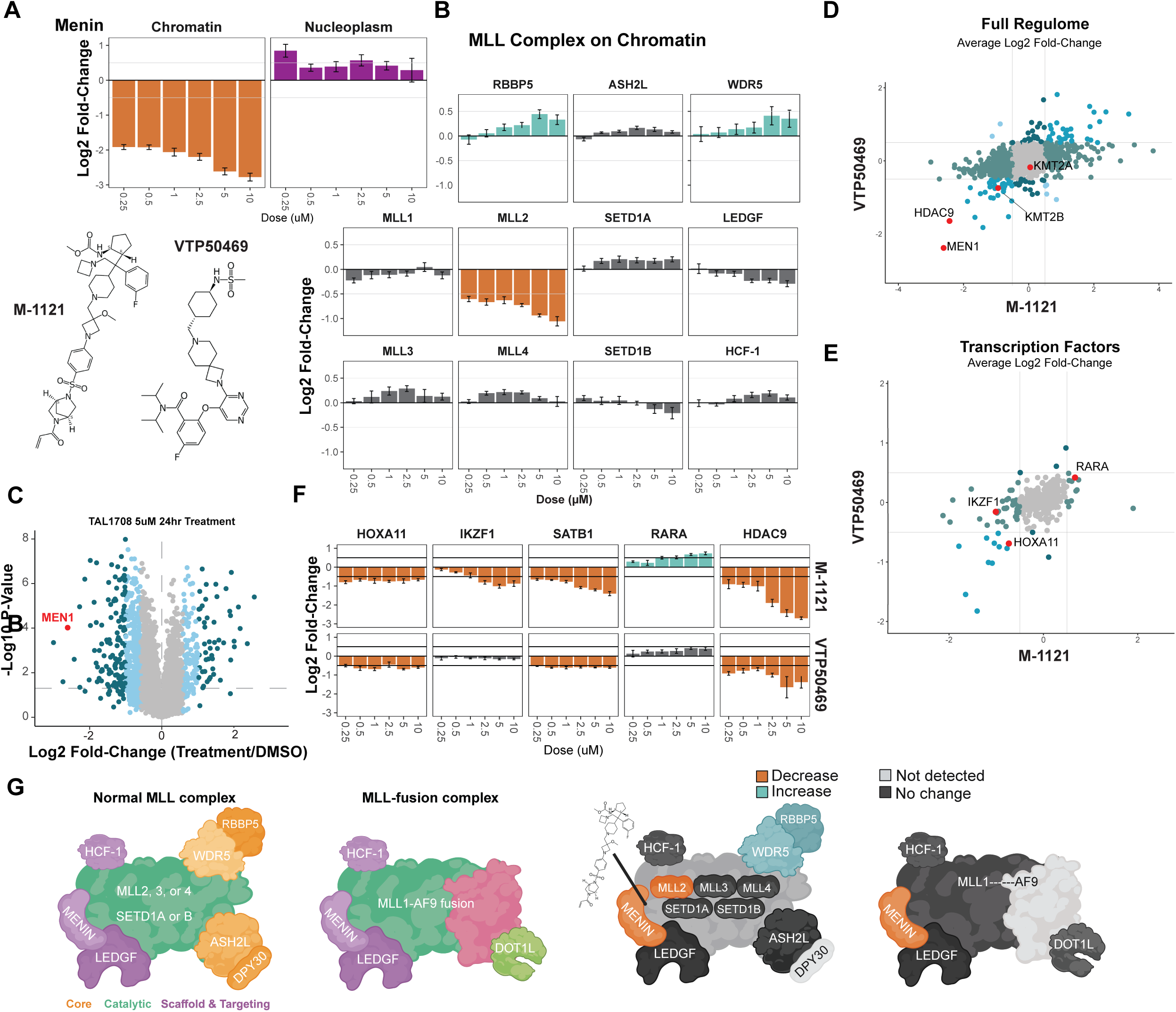
MLL-containing complex dynamics upon MLL-Menin inhibition (A) Menin levels measured across chromatin and nucleoplasmic fractions in THP-1 cells following treatment with M-1121 or VTP50469. Chromatin-associated Menin is depleted in a dose-dependent manner by both compounds, with a reciprocal increase in the nucleoplasmic pool. (B) Abundance of core MLL complex components in the chromatin fraction following Menin inhibitor treatment. (C) Volcano plot showing global protein-level changes in chromatin following treatment with M-1121 (5 µM, 24 h). (D) Comparison of average log₂ fold-changes across the entire regulome following treatment with M-1121 vs. VTP50469. (E) Subset of transcription factors from panel D, highlighting divergent effects on IKZF1, RARA, and HOXA11. (F) Dose-response curves for five selected TFs and cofactors impacted by Menin inhibition, with M-1121 and VTP50469 shown in parallel. (G) Schematic of canonical MLL complexes in wild-type and MLL-rearranged settings. Menin is essential for assembly and chromatin localization in the fusion complex, whereas core catalytic activity persists in the absence of Menin.

Outside the primary target, MLL2 was depleted from chromatin following treatment with both compounds, whereas other MLL paralogs remained stable. Core COMPASS components WDR5 and RBBP5 showed modest increases in chromatin association, which may reflect compensatory stabilization of alternative assemblies or changes in chromatin accessibility that promote retention of the scaffold (**Figure 5B**). Both compounds also triggered remodeling events beyond the MLL complex, including depletion of HDAC9 and HOXA11, with M-1121 specifically reducing IKZF1—a factor previously shown to co-occupy MLL fusion target loci with Menin and KMT2A (**Figure 5D–F**). Notably, RARA was modestly enriched in chromatin fractions following M-1121 treatment, suggesting potential priming for ATRA-induced differentiation in MLL-rearranged AML, consistent with prior studies (Sakamoto et al., 2014).

Together, these findings illustrate how covalent Menin inhibition perturbs not only the core MLL1 complex but rewires the broader regulatory proteome, revealing mechanistic nodes with potential therapeutic relevance (**Figure 5G**).

## DISCUSSION

Emerging models being developed towards the goal of the virtual cell rely on high-quality, multi-omic data to reconstruct and predict cellular behavior. Here we present a scalable approach for measuring the regulome directly at scale, with high quality, across cell types and perturbations.

By enriching for chromatin-associated proteins and quantifying them using label-free mass spectrometry, we generate a dataset that is complementary to RNA and whole-cell proteomics. The initial Regulome Atlas spans 36 human cell lines and recovers nearly 60 percent of known TFs, with subcellular resolution that distinguishes active chromatin-bound regulome proteins from sequestered regulome proteins in the cytoplasm and nucleoplasm. The method captures diverse families that participate in gene expression including TFs, cofactors, RNA binders, and chromatin-modifying complexes in an accessible, robust approach.

Several limitations of this warrant consideration. First, regulome profiling measures chromatin association rather than direct functional activity, and high abundance does not necessarily equate to transcriptional impact. particularly for proteins with varying specific activities, post-translational states, or competitive binding dynamics. However, many regulatory functions emerge not from single-factor activity, but from coordinated action among transcription factors, cofactors, chromatin remodelers, and RNA-binding proteins. As such, the relevant functional information may be embedded in the broader structure of the regulome itself. By capturing this context, regulome profiling may offer a more faithful reflection of transcriptional state than isolated measurements of any individual factor.

A second limitation is potential bias toward more abundant proteins due to the dynamic range limitations of mass spectrometry, potentially underrepresenting low-abundance but functionally important regulators. Despite these limitations, the strong concordance with established biological knowledge and orthogonal methods supports the biological relevance of these measurements.

Looking ahead, this approach enables broad discovery and application, thanks to its low sample input and experimental accessibility. Integrating regulome profiles with perturbation-response datasets could yield predictive models of transcriptional state transitions. Expanding the atlas to include primary tissues, developmental stages, and disease contexts would reveal regulatory dynamics not captured in cell lines. And combining regulome data with RNA-seq, chromatin accessibility, and metabolomics may offer a multilayered view of how cells process signals and control fate. In the long term, incorporating post-translational modifications or extending to single-cell resolution could uncover regulatory depth and heterogeneity that remain inaccessible today.

In summary, regulome profiles illuminate unbiased responses to diverse perturbations, capturing the downstream regulatory outcomes of cellular signaling that remain invisible to transcriptomic or proteomic profiling.. The method also can detect drug-target engagement and remodeling of chromatin-associated complexes during perturbation. These capabilities position regulome profiling as a powerful tool for decoding how cells interpret the environment and respond with genome regulatory programs, and as a foundational layer for building predictive models of cellular behavior.

## METHODS

### Cell culture and drug treatments

All cell lines were procured from ATCC and cultured as recommended. For compound and stimulant dosing, compounds were added using an Echo liquid handler to dispense nanoliter volumes of test compound or DMSO into each well. Doses were customized for each compound based on a literature review and are noted in Supplementary Table 2. After 24 hours, the cells were prepared for automated cell fractionation and regulome capture.

### Regulome purification

Cells were plated in 96 well plates at a seeding density specific to each cell type. Cells were treated with cytoplasm extraction buffer and co-incubated with lectin-coated magnetic beads at 4C for 20 minutes to immobilize intact nuclei. Immobilized nuclei were manipulated with a Kingfisher Flex maintained at 4C and exposed to a series of buffers to extract unbound nucleoplasm proteins, soluble chromatin proteins, leaving insoluble chromatin proteins attached to the bead. Protein fractions were prepared using beads and buffers contained in the Talus Bio Regulome Purification Kit for 96 well-plates (available after publication as TB1096, Talus Bio). Extracted proteins were stored at −80C until digestion.

### Preparation of proteins for analysis by mass spectrometry

Protein solutions were enzymatically digested by trypsin into peptides using an automated single-pot, solid-phase-enhanced sample preparation for proteomics (SP3) protocol ^32^. Briefly, proteins are denatured at 95C for 5 minutes, reduced with 4 mM DTT, alkylated with 20uM IAM, quenched with 4 uM DTT. Prepared proteins are then aggregated onto carboxyl beads (Cytiva) with 100% EtOH, washed using a Kingfisher Flex, incubated overnight at 37C with 1:10 trypsin:protein, and the resulting peptides prepared for LC-MS on Evotips (Evosep) using an OpenTrons liquid handler.

### Data independent acquisition mass spectrometry with Bruker timsTOF Ultra II

Peptides were analyzed with an Evosep One coupled with a Bruker timsTOF Ultra II mass spectrometer. We used a 8 cm x 150 µm “EV1109 Performance Column – 60/100 SPD”, a commercially available analytical column for the Evosep system with pre-mounted connection fittings packed with ReproSil Saphir C18, 1.5 µm beads by Dr Maisch. Solvent A was 0.1% formic acid in water, while solvent B was 0.1% formic acid in acetonitrile. For each injection, we loaded approximately 100 ng peptides and separated them using the Evosep 60 samples per day (SPD) method. Mass spectrometry diaPASEF measurement was acquired using 25 m/z isolation windows spanning a range from 400 to 1000 m/z and 8×3 TIMS ramps spaced from 0.64 to 1.45 1/K0. 75ms accumulation and 75 ramp cycles were used.

### Mass spectrometry data processing

Mass spectrometry data files were analyzed by DIA-NN (version 1.8.1) using the quantms Nextflow pipeline ^34^. Library free search was performed using a human FASTA from Uniprot including isoforms (accessed 2022-03-15) and supplemented with common contaminants. The following parameters were changed from default: mass_acc_automatic = false; diann_normalize = false.

## Data Availability

Raw mass spectrometry data will be made public in PRIDE upon acceptance. Processed regulome abundance matrices and analysis code are available upon request. We are developing an interactive data browser to facilitate community access to the Regulome Atlas and welcome collaborative opportunities to expand this resource.

## ACKNOWLEDGEMENTS

This work was supported in part by the National Institutes of Health (NIH 1R44TR004349, 4R44TR004349, 1R43TR004221, 1R42CA268146 to AJF; 1R43GM146472 to WEF), the National Science Foundation (NSF 2112191 to AJF), and the Andy Hill CARE Fund (FY23-LS-11 to AJF; FY23-LS-12 to LKP). We thank Noelle Schumacher, Bodhi Hueffmeier, Dr. Yang Gao, Dr. Tonibelle Gatbonton-Schwager, Dr. Charles Lin, Dr. Christopher Ott, Dr. Berkeley Gryder, Dr. Benedetta di Robilant, and Dr. Jun Qi for serving as early users of the platform and providing critical feedback during development. We are also grateful to all following for technical guidance and valuable discussions: Prof. John Stamatoyannopoulos, Prof. Michael J MacCoss, Prof. Brian C. Searle, Dr. Avery Sonnenberg, Dr. J Seth Stratton, Dr. Omri Amirav-Drory, Dr. Ronald Paranal, Mr. Seth Bannon, Dr. Bridget Martell, Dr. Michael McKeown, Dr. Stephen Letrent, Dr. Tara Arvedson, Dr. Andrei Konradi, Dr. Gavin Hirst. Margaux McBirney, Angelica Baca, and Michelle Briscoe provided essential operational support throughout the project.

